# Effect of partial substitution of rice with sorghum and inclusion of hydrolyzable tannins on digestibility and postprandial glycemia in adult dogs

**DOI:** 10.1101/482083

**Authors:** Liege Teixeira, Caroline Fredrich Dourado Pinto, Alexandre de Mello Kessler, Luciano Trevizan

**Author notes:** Corresponding author (LT).

## Abstract

Sorghum is used as a substitute of rice in dog food, owing to its nutritional similarity and low cost. However, its use has been associated with negative effects, like a reduction in palatability, digestibility, and enzyme activity, which can decrease nutrient absorption. The presence of condensed tannins (CT) in sorghum may cause these effects. Another tannin group, the hydrolysable tannins (HT), is known for its antioxidant properties. Research has shown the nutritional effects of sorghum on dogs, but the effect of HT on dogs remains unknown. We evaluated the effects of substituting rice with sorghum containing CT and inclusion of commercial extract of HT on digestibility, fecal and urinary characteristics, and postprandial blood glucose levels in adult dogs. Nine adult Beagle were randomly subjected to 4 treatments: 50% rice; (RS) 25% rice + 25% sorghum; (RHT) 50% rice + 0.10% HT; (RSHT) 25% rice + 25% sorghum + 0,10% HT. Tannins did not affect food intake. The digestibility of dry matter, organic matter, crude protein, acid hydrolyzed fat, gross energy, and metabolizable energy (ME) decreased with sorghum inclusion (P < 0.05). Sorghum also decreased protein digestibility (P < 0.05). Greater fecal dry matter was observed with the RHT diet. HT associated with sorghum reduced ME (P < 0.05). Sorghum inclusion enhanced fecal output, without altering fecal score (P > 0.05). No alterations in urinary characteristics were observed. Sorghum and HT did not affect the postprandial blood glucose response measured by the area under the curve (P > 0.05). The substitution of rice by sorghum negatively affected protein absorption and ME of the diets. Sorghum can be considered as a good source of carbohydrates in therapeutic diets for weight control. HT may potentiate the effect of CT, but more research is needed to evaluate its potential use in dog nutrition.

## Introduction

Carbohydrates are the main source of energy for most commercial dry-extruded diets for adult dogs, with cereal grains representing 30 - 60% of the final formula [1]. Rice is a functional ingredient regularly used in extruded dog food, due to its high digestibility. But, with the growing pet population over the past years, as well as the pet food industry, the search for alternative ingredients to provide nutritional quality and functional properties is becoming increasingly important [2].

In this scenario, sorghum (*Sorghum bicolor* L. Moench) appears as a viable option owing to its high productivity per hectare, drought tolerance, resistance to pests, good nutritional value, and lower cost of production, when compared with rice and corn [3]. Sorghum is commonly used—partially or wholly—as a source of energy in diets for non-ruminant animals, such as pigs and poultry [4,5]. It has been associated with some negative effects, however, especially on animal performance. Research has shown that those negative effects are linked to the presence of phenolic compounds, particularly tannins, which are secondary compounds of plant metabolism that affect different biological processes through their antimicrobial, antiparasitic, antioxidant, anti-inflammatory and antiviral properties [6]. However, tannins can also inhibit enzymes and form complexes with carbohydrates, proteins, and metal ions, thereby reducing nutrient intake and digestibility [7]. Tannins are classified according to their chemical structure into condensates (CT) and hydrolysables (HT). CT, designated as proanthocyanidins, are polymers of flavan-3-ols and flavan-3,4-diols, which can be oxidized to yield anthocyanidins. They are compounds resistant to hydrolysis, but soluble in aqueous organic solvents according to their chemical structure [8]. The HT are composed of simple phenols, gallotannins and elagitannins, which after hydrolysis produce gallic acid and ellagic acid. They are more easily hydrolyzed by acids and bases, and in some cases by enzymatic hydrolysis [9]. Sorghum is a solely source of CT [10]. HT are present in the leaves, flowers, twigs, and bark of some plants, and can also be found in a purified form as a commercial extract [11].

Although sorghum may contain varying levels of antinutritional factors, it is composed of up to 70% starch, of which 70–80% occur in the form of amylopectin and 20–30% occur as amylose [12]. Similarly, rice contains 75% starch, of which up to 35% is amylose [13,14]. The amylose / amylopectin ratio is one of the main indicators used to determine starch digestibility [15]. Amylopectin has a higher gelatinization capacity during the extrusion process, which increases starch digestibility. On the contrary, amylose possesses greater power of retrogradation during the same process, which reduces starch digestibility [16].

In addition to its effects on digestibility, starch is the main dietary component responsible for variation in postprandial glycemia in animals [17]. The faster and more complete the digestion, the faster and more intense glycemic the curve [18]. The slower digestion of amylose-rich starch appears to reduce the glycemic rate of animals by releasing glucose gradually into the bloodstream. Carciofi et al. [19] observed greater immediate postprandial glucose response for rice and corn and later response for sorghum in adult dogs.

Therefore, we hypothesized that sorghum can help modulate glycemic absorption due to the action of phenolic compounds. Thus, the ingredient is slowly digested, contributing to a longer satiety, delaying gastric emptying, and allowing slow glucose uptake compared to other cereals [10,20]. Such properties may be useful in cases of obesity, which - when untreated - can lead to decreased longevity, diabetes mellitus, orthopedic and respiratory diseases [21]. Some studies have observed a slight reduction in the digestibility of some nutrients with the inclusion of sorghum in the diets for dogs, although the supplemented diet was still accepted for commercial purposes [1,22]. Based on this evidence and on the lack of complementary information of the effect of sorghum on postprandial glycemia in adult dogs, this study aims to evaluate the partial replacement of rice by sorghum with CT and the inclusion of HT commercial extract, and their combined effect on the digestibility and postprandial glycemic response in adult dogs.

## Materials and methods

All animal care and handling procedures were approved by The Institutional Animal Care and Use Committee at the Federal University of Rio Grande do Sul, protocol number 26.275.

### Animals

Nine healthy adult Beagle (5 males and 4 females) were used in this study. They were all intact, between 2 and 3 years old, weighing 12.4 ± 0.97 kg, with a body condition score (BCS) ranging from 4.5 to 5.5 out of 9 points [23] and free of endo- and ectoparasites. All dogs were regularly immunized and submitted to clinical and laboratory tests to measure complete blood count (CBC) and to perform biochemical and coproparasitological analyses before the start of the study. The dogs were housed in individual stainless steel metabolic cages (1.0 × 1.0 × 1.5 m) equipped with a feces and urine collector, feeders, and drinkers, in a controlled room at 24°C, with a light:dark cycle of 14:10 h.

### Diets

Rice was partially substituted with sorghum as a way to introduce CT into the diets. Additionally, purified HT obtained from a commercial extract of the chestnut bark (Silvafeed ENC^®^, Piedmont, Italy) was included into the diets. The extract was obtained by heating the chestnut bark with water at low pressure, and subsequently dehydrating the water-soluble fraction. The final product was a fine brown powder containing HT, hydrolyzable polyphenols, cellulose, hemicellulose, simple sugars, lignin, minerals, and 8% moisture; its fiber content was < 3% and it had a relative density of 0.5–0.6% and pH < 4.0. Four experimental diets were formulated and extruded to be isonutritives: (R) 50% rice; (RS) 25% rice + 25% sorghum; (RHT) 50% rice + 0.10% HT; (RSHT) 25% rice + 25% sorghum + 0.10% HT (see Table 1 and 2). The dogs were fed twice a day (at 8:30 and 17:00) to meet the energetic and nutritional requirements of adult dogs, as recommended by the NRC [24]. The leftovers were collected, weighed, and discounted to calculate consumption. Water was provided *ad libitum*.

**Table 1.** Chemical composition of sorghum (*Sorghum bicolor* L. Moench)

| Item, % DM basis | Sorghum |
| --- | --- |
| DM | 86.9 |
| Starch | 63.6 |
| Crude protein | 7.59 |
| Ether extract | 2.56 |
| Ash | 1.42 |
| Crude fiber | 0.72 |
| Gross energy, kcal/kg | 4.446 |
| Polyphenol tannins, % | 4.8 |
| Polyphenol non-tannins, % | 2.6 |

DM, dry matter.

**Table 2.** Ingredients and chemical composition of experimental diets

| Ingredient, % | Treatments |
| --- | --- |

|  | R | RS | RHT | RSHT |
| --- | --- | --- | --- | --- |
| Broken rice | 50.7 | 26.2 | 50.7 | 26.2 |
| Sorghum | - | 25.0 | - | 25.0 |
| Hydrolysable tannins <sup>1</sup> | - | - | 0.10 | 0.10 |
| Wheat bran | 14.0 | 14.0 | 14.0 | 14.0 |
| Poultry byproducts meal | 11.1 | 11.4 | 11.1 | 11.4 |
| Meat and bone bovine meal | 8.00 | 8.00 | 8.00 | 8.00 |
| Poultry fat | 6.00 | 5.81 | 6.00 | 5.81 |
| Corn gluten 60% CP | 5.00 | 5.00 | 5.00 | 5.00 |
| Digest <sup>2</sup> | 1.50 | 1.50 | 1.50 | 1.50 |
| Cellulose | 1.17 | 1.15 | 1.17 | 1.15 |
| Flaxseed | 1.00 | 1.00 | 1.00 | 1.00 |
| Soybean oil | 0.52 | - | 0.52 | - |
| Premix mineral/vitamin <sup>3</sup> | 0.40 | 0.40 | 0.40 | 0.40 |
| Salt | 0.38 | 0.38 | 0.38 | 0.38 |
| Potassium chloride | 0.17 | 0.07 | 0.17 | 0.07 |
| Starch | 0.10 | 0.10 | 0.00 | 0.00 |
| Total | 100 | 100 | 100 | 100 |
| Analyzed chemical composition, % DM basis |  |  |  |  |
| Dry matter | 91.6 | 87.0 | 88.2 | 90.3 |
| Starch | 39.6 | 38.0 | 39.6 | 37.7 |
| Crude protein | 19.5 | 20.7 | 21.5 | 18.8 |
| Acid hydrolyzed fat | 8.99 | 8.71 | 9.05 | 8.80 |
| Ash | 6.34 | 7.11 | 7.54 | 7.39 |
| Crude fiber | 3.64 | 4.21 | 3.64 | 4.13 |
| Total dietary fiber | 22.5 | 21.2 | 22.5 | 22.7 |
| GE, kcal/kg | 4.903 | 4.881 | 4.799 | 4.815 |
| Gelatinization index of starch, % | 92.0 | 91.3 | 90.2 | 92.3 |
| Polyphenol tannins, % | - | 1.2 | 0.1 | 1.3 |

R, rice; RS, rice + sorghum; RHT, rice + hydrolysable tannins; RSHT, rice + sorghum + hydrolysable tannins; CP, crude protein; GE, gross energy.
<sup>1</sup> Silvaeed ENC<sup>®</sup>, Piedmont, Italy.
<sup>2</sup> DTECH 8L, S.P.F. Argentina S.A., Argentina.
<sup>3</sup> Premix composition per kilogram: vitamin A (10.800 UI), vitamin D3 (980 UI), vitamin E (60 mg), vitamin K3 (4.8 mg), vitamin B1 (8.1 mg), vitamin B2 (6.0 mg), vitamin B6 (6.0 mg), 12 vitamin (30 mcg), pantothenic acid (12 mg), niacin (60 mg), folic acid (0.8 mg), biotin (0.084 mg), manganese (7.5 mg), zinc (100 mg), iron (35 mg), copper (7.0 mg), cobalt (10 mg), iodine (1.5 mg), selenium (0.36 mg), choline (2.400 mg), taurine (100 mg), and, antioxidant BHT (150 mg).

## Experiment 1: Digestibility Assay

### Experimental Design

The assay was conducted as an incomplete randomized block design of four treatments and three 10-day periods, with 2 dogs per treatment in each period, for a total of 6 replications per treatment, according to the recommendations of the American Association of Feed Control Officials protocol [25]. Each period lasted 10 days, with 5 days for adaptation to the cage and experimental diet, followed by 5 days of total feces and urine collection and measurement of fecal and urinary pH. Between each period 5 days of rest were provided to the dogs so they could exercise in which the R diet was provided.

### Sample procedure

To establish the beginning and the end of each period of feces and urine collection, gelatin capsules containing 1 g of iron oxide (III) Fe_2_O_3_ were orally given to the dogs. Feces were collected for 5 days and scored as follows: 1 = very hard and dry stool, 2 = hard, dry, firm stool, 3 = soft, moist stool, well formed, 4 = soft and shapeless stool, 5 = liquid stool, diarrhea. After daily collection, feces were weighed and stored in a freezer at - 20°C until the end of the trial to perform analysis. Total urine collection was performed daily in the morning and then stored in plastic bottles containing 1 g of thimol (Synth, Diadema, Brazil) and the pH was measured. The urine total volume was measured and kept in a freezer at - 20°C until analysis. The fecal pH was measured in 2 g of fresh feces diluted in 20 mL of distilled water (Digimed DM-22, Campo Grande, Brazil).

### Chemical Analysis

Stool from each dog was thawed, homogenized, and dried in forced-air oven at 55°C for 72 h, according to the recommendations of the Association of Official Analytical Chemists [26]. Feces and diets were ground through a 1 mm screen in a Wiley hammer mill (DeLeo Equipamentos Laboratoriais, Porto Alegre, Brazil), and analyzed for dry matter (DM - AOAC 934.01), acid hydrolyzed fat (AOAC 954.02; model 170/3, Fanem, São Paulo, Brazil), crude protein (CP - AOAC 954.01; model TE 036/2, Tecnal, Piracicaba, Brazil), crude fiber (CF - AOAC 962.10; model MA 450/8, Marconi, Piracicaba, Brazil), and ash (MM) [26]. Urine samples were thawed, homogenized and 150 mL aliquots were lyophilized (Micromodulyi-Fis; Termo Fisher Scientifics INC, Maryland, USA) for analysis of DM and gross energy (GE). Another 50 mL aliquot was collected for analysis of CP. Dietary, fecal, and urinary GE were determined using isoperibolic bomb calorimetry (calorimeter model C2000 basic, Ika-werke, Staufen, Germany). All analyses were performed in duplicates, assuming a variation <1% for energy and <5% for the other analyzes. The tannins were analyzed by gravimetric tests using the method of Freiberg-Hide [27].

### Statistical Analyses

Data were analyzed using the ANOVA procedure of SAS 9.4 (SAS Inst. Inc., Cary, NC). Means were compared using Tukey’s test at 5% probability (*P* < 0.005).

## Experiment 2: Postprandial Glycemia

The dogs and the dietary treatments were the same as previously described for the digestibility assay.

### Experimental Design

The dogs were adapted to the experimental diets for 11 days, then were fasted for 12 h inside the metabolic cages before starting the first blood collection. Immediately before starting the experiment, the cephalic vein was cannulated with a catheter BD ANGIOCATH® 22’’ (Becton, Dickinson and Company do Brasil, Curitiba, Brazil). Then, 1 mL of blood was collected in a tube containing 0.05 mL of sodium fluoride (LABTEST®, Lagoa Santa, Brazil); this sample was used to determine the baseline glycemia at time 0. Then, food was offered and was consumed in 5 min by all the dogs. Sequential collections were started over 8 h, at 5, 10, 15, 30, 45, 60, 90, 120, 180, 240, 300, 360, 420, and 480 min after food consumption. After each collection the catheter was washed with heparinized solution and before each new collection, about 0.3 mL of blood were discarded.

### Chemical Analyses

The tubes were centrifuged at 3000 *g* during 10 min, and plasma was transferred to Eppendorf tubes of 1.5 mL, cooled between 2 and 4°C, and analyzed in sequence. Blood glucose was analyzed by the enzymatic colorimetric method according to the manufacturer’s instructions (Wiener Lab Group, Rosário, Argentina). All samples were analyzed in duplicate.

### Statistical Analyses

The results were analyzed using the ANOVA with repeated measures in time. The area under the curve (AUC) was calculated, and the mean of each treatment was compared by Tukey’s test (*P* < 0.05) using the SAS 9.4 (SAS Inst. Inc., Cary, NC).

## Results

The dogs normally consumed all the experimental diets offered quickly, without refusal. The inclusion of sorghum and HT did not promote clinical alterations such as vomiting and diarrhea. Initial and final CBC and biochemical profiles remained within the normal range for adult dogs [28].

Sorghum contained around 4.8% of polyphenol tannins (see Table 1), and was the only ingredient in the diets containing a significant amount of tannins. According to this inclusion, 1.2% of tannins compound is ensured in the diets. The food was well cooked, based on the gelatinization of starch, and all the diets presented a gelatinization index greater than 90%. Diets had small differences in nutrient concentration owing to rice substitution with sorghum (Table 2), which had some influence on the variable nutrient uptake for the different diets (see Table 3). The intake of crude fiber was higher in dogs fed diets containing sorghum (*P* < 0.0005) but this was not the case when total dietary fiber was evaluated, as dogs had the same dry matter consumption, and the concentration of total dietary fiber was similar among diets. Dogs fed diets containing CT and HT together consumed more ash (*P* < 0.0164).

**Table 3.** Nutrient intake, and apparent total tract digestibility of macronutrients and energy of dogs fed experimental diets

| Item | Diets |  |  |  | *P*-value | SEM |
| --- | --- | --- | --- | --- | --- | --- |
|  | R | RS | RHT | RSHT |  |  |
| Daily nutrient intake, g |  |  |  |  |  |  |
| DM | 206 | 201 | 194 | 197 | 0.4223 | 12.7 |
| OM | 193 | 187 | 179 | 183 | 0.2635 | 11.8 |
| AHF | 18.5 | 17.5 | 17.6 | 17.4 | 0.3254 | 1.13 |
| CP | 40.0<sup>a</sup> | 41.7<sup>a</sup> | 41.8<sup>a</sup> | 36.8<sup>ab</sup> | 0.0136 | 2.42 |
| FDT | 37.5<sup>b</sup> | 42.3<sup>a</sup> | 35.3<sup>b</sup> | 40.7<sup>ab</sup> | 0.0005 | 2.43 |
| Ash | 13.0<sup>b</sup> | 14.3<sup>ab</sup> | 14.6<sup>a</sup> | 14.6<sup>a</sup> | 0.0164 | 0.85 |
| NFE | 134<sup>a</sup> | 128<sup>ab</sup> | 120<sup>b</sup> | 124<sup>ab</sup> | 0.0496 | 8.21 |
| Apparent total tract digestibility, % |  |  |  |  |  |  |
| DM | 81.8<sup>a</sup> | 78.5<sup>ab</sup> | 81.2<sup>a</sup> | 76.7<sup>b</sup> | 0.0015 | 1.95 |
| OM | 85.3<sup>a</sup> | 81.9<sup>b</sup> | 85.3<sup>a</sup> | 80.9<sup>b</sup> | 0.0007 | 1.74 |
| AHF | 92.8 | 89.4 | 92.6 | 90.3 | 0.2122 | 3.06 |
| CP | 84.9<sup>a</sup> | 82.3<sup>ab</sup> | 87.0<sup>a</sup> | 79.6<sup>b</sup> | <0.0001 | 1.94 |
| CF | 31.7 | 30.8 | 37.2 | 33.7 | 0.6901 | 10.1 |
| Ash | 30.2 | 33.2 | 30.8 | 24.3 | 0.2224 | 6.93 |
| NFE | 84.4<sup>a</sup> | 80.8<sup>bc</sup> | 83.6<sup>ab</sup> | 79.3<sup>c</sup> | 0.0015 | 1.96 |
| GE | 85.4<sup>a</sup> | 82.1<sup>b</sup> | 85.2<sup>a</sup> | 80.7<sup>b</sup> | 0.0004 | 1.69 |

| Nutricional value, kcal/kg |  |  |  |  |  |  |
| --- | --- | --- | --- | --- | --- | --- |
| ME | 3.978<sup>a</sup> | 3.793<sup>bc</sup> | 3.857<sup>ab</sup> | 3.698<sup>c</sup> | 0.0002 | 78.7 |

R, rice; RS, rice + sorghum; RHT, rice + hydrolysable tannins; RSHT, rice + sorghum + hydrolysable tannins; SEM, standard error of the mean; DM, dry matter; OM, organic matter; AHF, acid hydrolyzed fat; CP, crude protein; CF, crude fiber; NFE, nitrogen-free extractive; GE, gross energy; ME, metabolizable energy.

Sorghum inclusion reduced the digestibility coefficients of DM, OM, CP, DE, and ME (*P* < 0.05), especially in the RSHT treatment (see Table 3), which contained both tannins. The inclusion of HT was not enough to reduce the ME content, only when added with sorghum (*P* < 0.0002). The nutrient and energy digestibility coefficients of the RHT treatment did not differ significantly from the control group (R), which presented the best results regarding digestibility compared to the others. The results suggest that there was a potentiate effect of tannins, CT plus HT, influencing the reduction in nutrient and energy digestibility.

The inclusion of HT reduced the fecal water content, and this effect was greater in dogs fed diets containing sorghum (*P* < 0.0045) (see Table 4). Dogs fed RHT treatment had lower daily fecal production compared to those treated with sorghum (*P* < 0.0059). Despite these alterations, the mean fecal score did not differ between diets, resulting in dry and firm stools, a desired aspect in the extruded diets.

**Table 4.** Fecal and urinary characteristics of dogs fed diets containing tannins

| Item | Diets |  |  |  | *P*-value | SEM |
| --- | --- | --- | --- | --- | --- | --- |
|  | R | RS | RHT | RSHT |  |  |
| Fecal characteristics |  |  |  |  |  |  |
| Fecal DM, % | 38.9<sup>ab</sup> | 38.4<sup>b</sup> | 41.1<sup>a</sup> | 37.6<sup>b</sup> | 0.0045 | 1.42 |
| Fecal output, g/d | 96.5<sup>b</sup> | 113<sup>ab</sup> | 88.6<sup>b</sup> | 123<sup>a</sup> | 0.0059 | 15.0 |
| Fecal output, g/d<br>(DM) | 37.4<sup>b</sup> | 43.2<sup>ab</sup> | 36.5<sup>b</sup> | 46.2<sup>a</sup> | 0.0107 | 4.83 |
| Feces pH | 6.65 | 6.60 | 6.70 | 6.65 | 0.7919 | 0.17 |
| Fecal score, 1 to 5 | 2.05 | 2.13 | 2.04 | 2.11 | 0.2050 | 0.08 |
| Urinary characteristics |  |  |  |  |  |  |
| Volume, mL<sup>1</sup> | 2334 | 2461 | 1663 | 2549 | 0.2782 | 835 |
| Total DM, % | 3.70 | 4.21 | 4.08 | 3.96 | 0.9547 | 1.66 |
| Urine CP, DM% | 7.75 | 8.78 | 9.46 | 8.52 | 0.8305 | 3.28 |
| Urine pH | 7.42 | 7.35 | 7.50 | 7.42 | 0.6975 | 0.21 |

R, rice; RS, rice + sorghum; RHT, rice + hydrolysable tannins; RSHT, rice + sorghum + hydrolysable tannins; SEM, standard error of the mean; DM, dry matter; CP, crude protein.
<sup>ab</sup>Letters superscript indicate the differences among diets.
<sup>1</sup>Volume produced in 5 days.

No change was observed in the urinary characteristics analyzed (*P* > 0.05) (see Table 4). However, the urine and feces produced by the dogs fed with diets containing sorghum presented a darker coloration than in the control diet (R).

The postprandial glycemic response, measured by the AUC, from 0 to 480 minutes after meal was not significantly different among groups. However, when the initial period is discounted, between 30 and 300 min, dogs fed the RHT diet tended to show the largest area under the curve (*P* = 0.07), meaning that absorption was greater than other diets in this period. But neither sorghum nor HT in the diets affected the basal, average, minimum, or maximum glycemia (*P* > 0.05) (see Table 5).

**Table 5.** Area under curve without basal glycemic area (AUC), plasma basal glucose concentration PBGC), plasma glucose concentration (PGC), and values of the glycemic peak

| Item | Diets |  |  |  | *P*-value | SEM |
| --- | --- | --- | --- | --- | --- | --- |
|  | R | RS | RHT | RSHT |  |  |
| AUC total (0-480) mg/dL*min | 38554 | 40350 | 41910 | 39615 | 0.35 | 705 |
| AUC (30-300) mg/dL*min | 16209 | 17182 | 18564 | 15936 | 0.07 | 1303 |
| Basal glycemia, mg/dL | 79.6 | 78.2 | 80.7 | 80.0 | 0.88 | 0.53 |
| Maximum glycemia, mg/dL | 92.6 | 99.7 | 102 | 97.2 | 0.42 | 2.01 |
| Average glycemia, mg/dL | 79.8 | 83.1 | 85.9 | 82.2 | 0.34 | 1.26 |
| Minimum glycemia, mg/dL | 71.6 | 68.7 | 74.1 | 71.2 | 0.44 | 1.11 |
| Maximum glycemic increase, mg/dL | 13.0 | 21.0 | 21.5 | 17.2 | 0.56 | 1.97 |

R, rice; RS, rice + sorghum; RHT, rice + hydrolysable tannins; RSHT, rice + sorghum + hydrolysable tannins; SEM, standard error of the mean; AUC, area under the curve.
Peak glycemia time of dogs fed experimental diet.

## Discussion

The objective of the present study was to determine if the inclusion of sorghum and HT in the diet would modify the digestibility of those diets, and how it would impact the postprandial glycemic index of adult dogs, based on the ability of tannins to form complexes with proteins and carbohydrates. Tannins are responsible for reducing enzyme activity in the gut, reducing the postprandial peaks of glycemia, promoting a long and flat curve of glycemia after the meal, and increasing the time of glucose absorption, as reported by Carciofi et al. [19]. In this study, we expected that sorghum would be used as a source of CT in canine diets, as rice and other sources of ingredients do not contain tannins. We hypothesized that sorghum or HT would work as functional ingredients to change the glycemic index of rice, which is considered to be an ingredient with high glycemic index. Diets were made to replace the included rice content by half and to maintain the same amount of starch and other macronutrients, including total dietary fiber content. HT was included to check for additional effect, as it has been used in diets for poultry and swine with some effects on feces quality. Schiavone et al. [29] used levels of 0.15%, 0.20%, and 0.25% of the same HT extract in broiler diets and did not obtain significant results in terms of digestibility and carcass quality related to the control diet. However, there was an increase in the DM content of the feces, and feces had a firmer consistency and a reduction in the water content. The presence of tannins did not alter the consumption of the diets: all dogs consumed quickly all the food provided. Acceptance is a crucial point when a new ingredient is tested. The diets formulated with sorghum and tannins were well acceptable by the dogs and the palatability test was not performed. There is no consensus on the effect of tannins on voluntary consumption, but it is known that the formation of complexes between tannins, proline-rich salivary proteins, and the mucosal epithelium of the oral cavity, and the direct connection of tannins with gustatory receptors, both contribute to the formation of tannins’ astringency [30]. Mole et al. [31] observed that dogs as well as cats produced small amounts of proline-rich salivary proteins, and did not have tannin affinity *in vitro*. This fact may partially explain the voluntary consumption we observed, as there was no precipitation of the complex tannin-proline-rich salivary proteins, thus no sensation of astringency. Similar results were observed in a study by Kore et al. [32], where dogs normally consumed the diet containing sorghum, without refusing or leaving leftovers.

Despite our attempt to make isonutritive diets, the substitution of rice with sorghum produced diets with some significant differences on nutrient consumption due to moisture of the final diets. The diets were mixed and extruded separately, which may have exacerbated this impact. The inclusion of sorghum negatively affected the digestibility of DM, organic matter (OM), CP, digestible energy, and ME of the diets. This fact may be due to the greater inhibitory power of CT on enzymatic activity compared to that of HT [33]. Kore et al. (2009) found similar results when evaluating different cereals for dogs, with sorghum presenting the lowest coefficients of digestibility of DM, OM, CP, and CF.

One of the main factors affecting the DM and OM digestibility coefficients is the starch gelatinization index. Starch grains absorb water, swell, release part of the amylose, and become more susceptible to enzymatic degradation [34]. The high gelatinization of the starch determines better extrusion and a better granule [35]. In this study, the gelatinization index was above 90% for all diets (see Table 2), indicating that the extrusion was effective to allow enzymatic access during digestion. Some researchers suggest that, during cooking, CT undergo structural modification through depolymerization into oligomers and monomers, but maintain the stable basic structure [3]. There are indications that in the intestine, the tannins will polymerize again and bind to proteins, forming compounds resistant to digestion. Thus, they reach the colon and may or may not be degraded into simple phenols [36]. The undigested tannins remain in the lumen, where they can antagonize the effects of pro-oxidants produced during the metabolism of microbiota [37]. Saura-Calixto et al. [38] did not observe any effect of digestive enzymes on the release and bioaccessibility of *in vitro* CT polymers in human diets, suggesting that they come unchanged in the colon. Another *in vitro* study using highly polymerized CT indicated that they were not affected by the intestinal microbiota [39].

The inclusion of HT did not affect the digestibility of the CP and did not differ significantly from the control treatment (R). There is some indication that HT must interact with proteins, forming less stable bonds than CT and allowing the intestinal microbiota to metabolize its components and make them more soluble [40]. Thus, HT may have a lower impact on digestibility than do CT. HT are absorbed mainly in the small intestine, being fermented in less quantity in the colon [39]. Hagerman et al. [41] observed a reduction in the digestibility of CP in sheep fed a diet containing CT, HT did not present the same effects. The reduction in DE and ME of the diets containing sorghum observed in this study concurs with previous studies [42]. DE and ME tend to decrease with the increase of dietary tannin content, through the formation of complexes with carbohydrates, reducing the activity of the amylolytic enzymes and their energetic use [43].

The addition of HT reduced the water content of dog feces, a desired trait in extruded diet. However, such an effect was not observed with the dietary association of HT and CT. The excess of non-digestible content into the lumen may have some impact on fecal water content. The dogs that consumed diets containing sorghum presented greater production of feces, due to the lower digestibility of the ingredient. However, the mean fecal score was not altered between diets, resulting in dry and firm stools. Twomey et al. [42] observed that dogs fed diets with sorghum produced firmer feces than those fed rice-based diets (*P* < 0.05), but all values were still within the ideal range.

No changes were observed in fecal and urinary pH. Dows et al. [44], using diets with CT in wild rodents, observed the production of more alkaline urine, which did not occur in this study. However, the urine and feces of dogs fed diets containing sorghum and HT presented a darker coloration than did the urine and feces of dogs fed rice-based diet, indicating the metabolism of tannins and excretion of their components in urine and feces. Purified tannins have darker coloration, which ranges from dark brown to black. Additionally, the sorghum used had a dark red coloration.

The presence of HT tended to promote a increase in the postprandial glycemic response of dogs, from 30 to 300 min, the time during which most parts of glucose are absorbed during the digestion. It was against our hypothesis, as we expected a reduction in the area under the curve. Hydrolysable tannins contain glucose in their molecular structure, to which galo- and elagio-tannin remains associated. From the dark color of the urine, it is possible to speculate that HT, including glucose present in HT, were digested; however, this is still unlikely to have produced an increase in glycemia, since the inclusion of HT in the diet was very less.

Carciofi et al. [19] observed a greater area under the curve for dogs fed sorghum compared to those fed with rice 30 min after the consumption of the experimental diets. The contrasting results can be explained by the variation in the chemical composition of the sorghum, which is influenced by genetic and environmental factors [45].

The amount of CT in sorghum has been reduced through genetic improvement of cultivars, allowing the grain to be used to feed non-ruminant animals without compromising digestibility and, consequently, animal performance. To test sorghum in the experimental diets we analyzed three varieties of sorghum to select the one with a higher concentration of condensed tannins.

Myer et al. [46] evaluated sorghum with different tannin levels for growing-finishing swine and considered a tannin content of 1.3 to 3.6% as high and 0.1 to 0.7% as low. The variety used in this study had a tannin content of 4.8%, classified as a high tannin grain sorghum. Although sorghum has no influence on the reduction of the postprandial glycemic response, the ingredient has desirable characteristics in specific products, such as calorie-restricted diets. Finally, more studies are needed to determine the actual effects of HT on dog health.

### Conclusions

The inclusion of tannins in canine diets did not affect the voluntary consumption of food by dogs. However, the presence of sorghum caused a reduction in the digestibility of DM, OM, CP, DE, and ME, and promoted greater fecal production. The fecal score was kept in good standard. There was a darkening of feces and urine of the dogs that received sorghum and HT in the diet, strongly indicating that there is metabolism and excretion of its constituents. The addition of HT reduced DM and water content in feces. Although the expected glycemic response results were not observed, sorghum-containing tannins have good applicability in canine diets. By reducing the ME content of the diets, sorghum can become a key ingredient in the development of therapeutic diets for weight control in dogs.

## Acknowledgement

The authors are thankful for the financial support given by Brazilian governmental research support institution Coordenação de Aperfeiçoamento de Pessoal de Nível Superior-CAPES and Nutribarrasul, Brasil, for the diets extrusion.

